## Supplementary TableS2 and figures S1-6 for "Chromosomal neighbourhoods allowidentification of organ specific changesin gene expression"

### Chromosomal neighbourhoods having genes of diverse functions allow improved identification of differentially expressed genes

Rishi Das Roy, Outi Hallikas, Mona M. Christensen, Elodie Renvoisé, Jukka Jernvall

#### Supplementary figures S1-6 and supplementary table S2

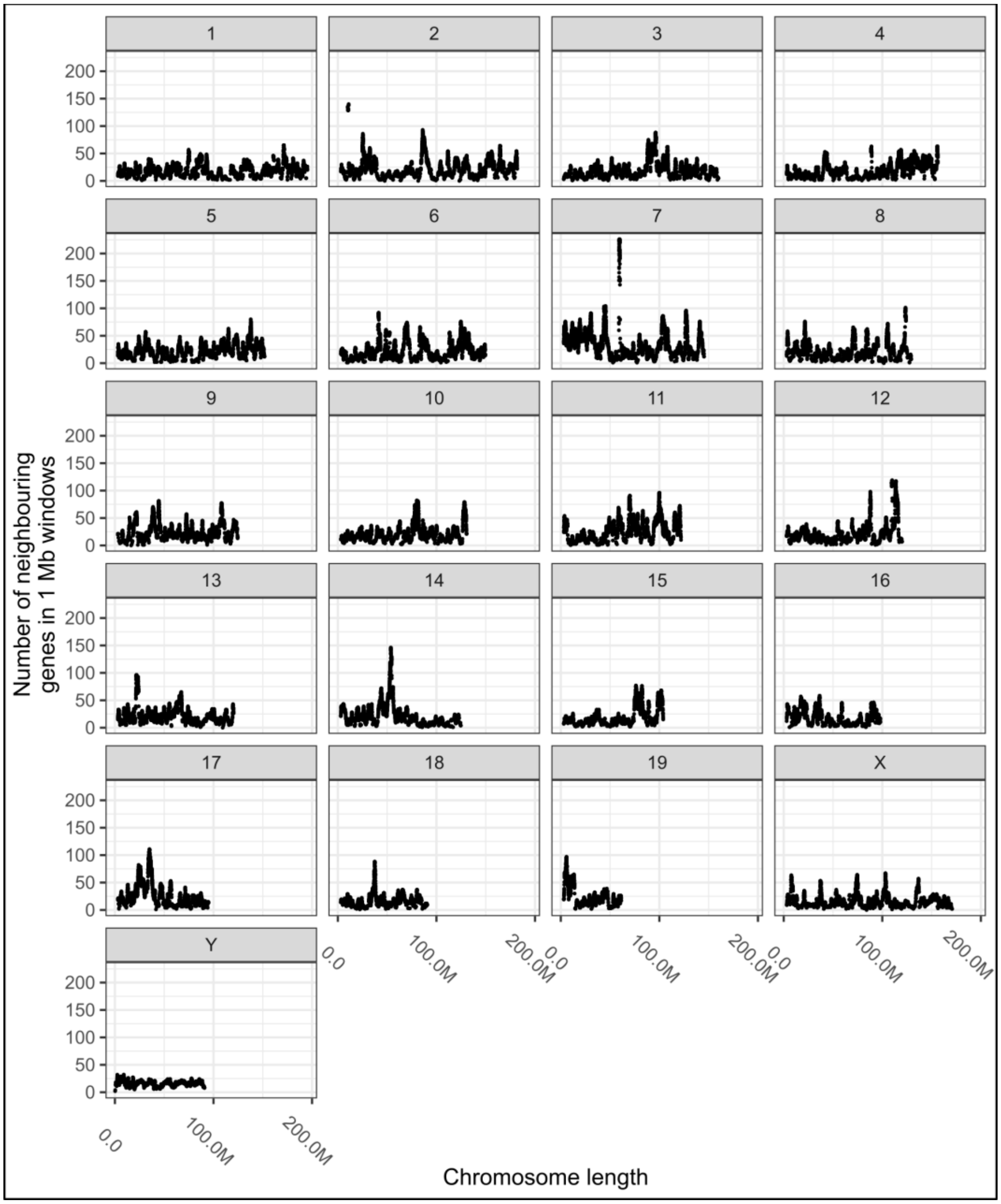


**Supplementary Figure S1**: Distribution of genes across mouse chromosomes (x-axis) and number of "neighbouring genes" (y-axis) within 1 Mb window around each gene.

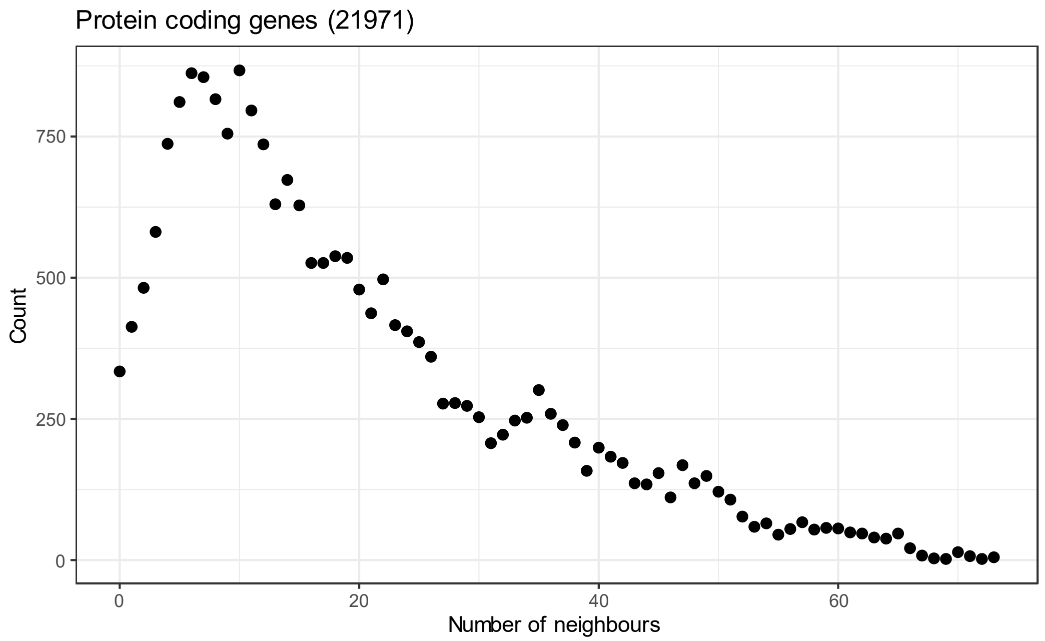


**Supplementary Figure S2:** Genes with 5-9 neighbouring genes are most frequent in the mouse genome. Each point represents the number of genes with the same number of neighbouring protein coding genes in 1 Mb window.

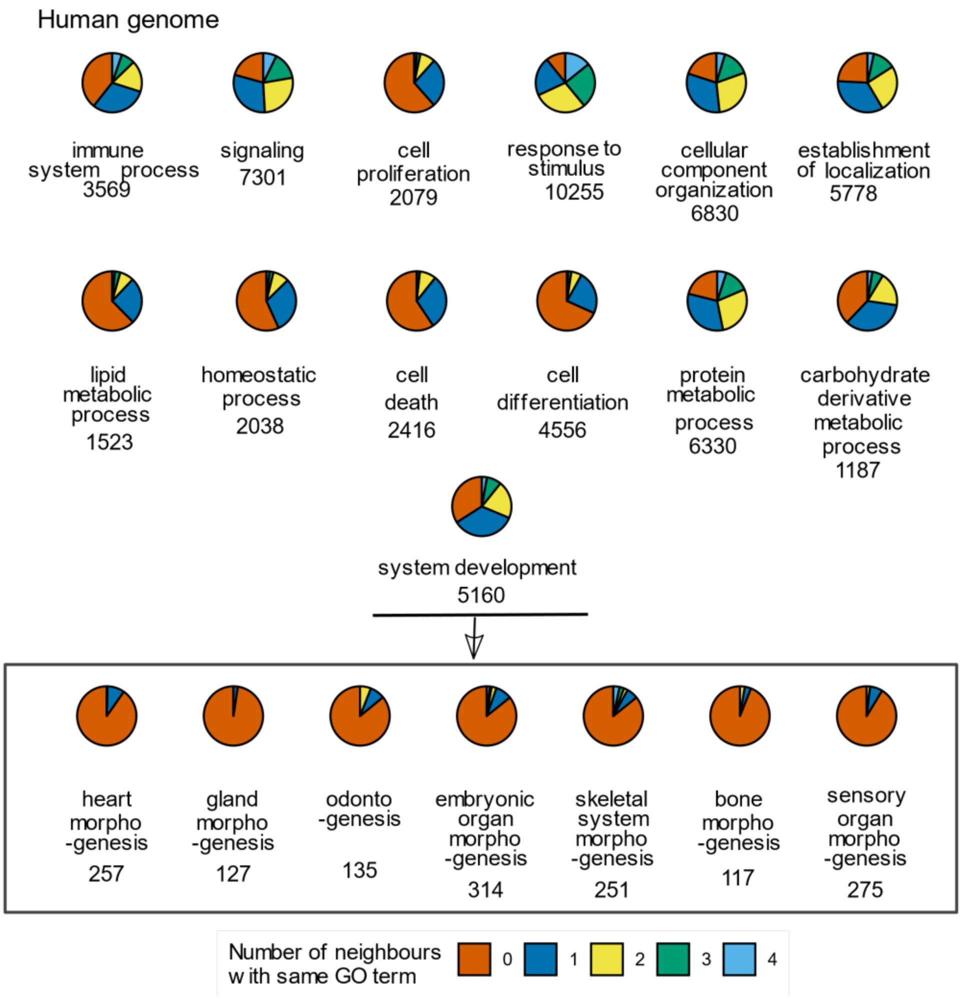


**Supplementary Figure S3.** The more specific the GO term, the fewer neighbours its genes have. Each pie represents the genes of a GO term under the root GO term 'biological processes'. The GO terms are arranged from top to bottom following their proximity to the root term 'biological process'. The color coding indicates the number of neighbours that a gene has from the same GO term. Analysis was done for the human genome. See Supplementary Table S2 for GO IDs.

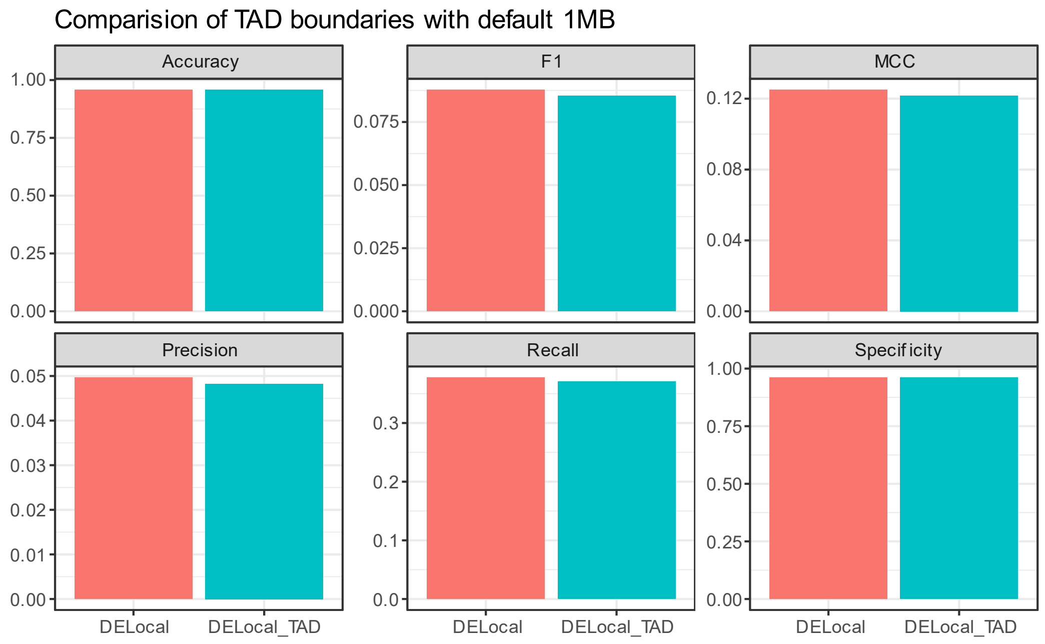


**Supplementary Figure S4.** Performance comparisons of DELocal calculated using neighbourhoods with 1 Mb windows and with TAD boundaries (from RNAseq data).

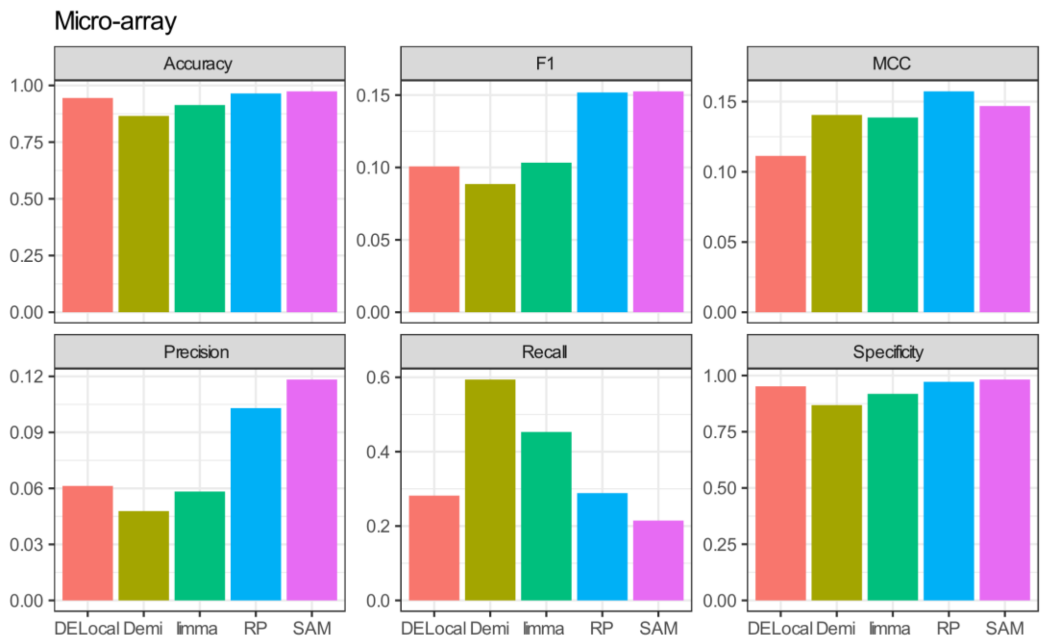


**Supplementary Figure S5**. Comparison of DELocal with earlier methods to identify differential expression. Evaluation matrices for microarray data. DELocal has lower performance in microarray data compared to RNAseq data (Figure 7), likely to be due to limited number of genes and neighbourhood data in microarrays. The evaluation matrices are explained in materials and method section.

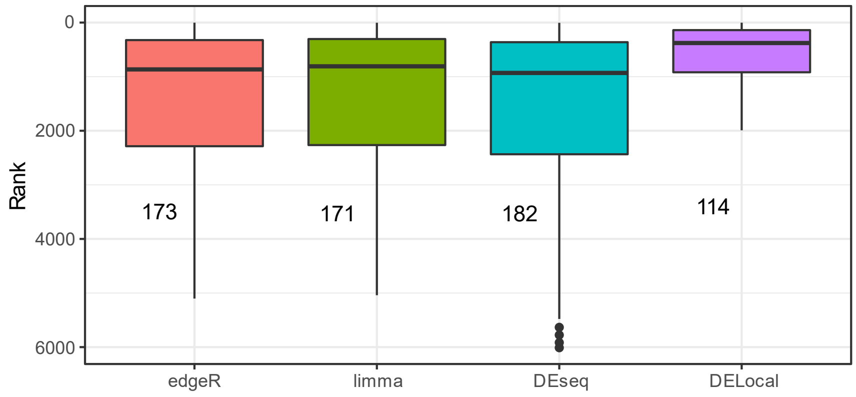


**Supplementary Figure S6.** Distribution of ranks of tooth developmental genes. DELocal predicted tooth developmental genes are significantly (Wilcoxon rank test; p-value <= 1.0e-06 ) enriched in top ranked positions compared to the other three methods. Although other methods identified more of the genes (numbers listed next to the box plots), this improved recall is sacrificing specificity. DELocal balances both which is also reflected in F1, MCC and ROC curves (Figures 6 and 7).

**Supplementary Table S2.** List of GO terms analysed in Figure 7,8 and S4.

| Display ID | GO ID | Description |
|---|---|---|
| 1 | GO:0060349 | bone morphogenesis |
| 2 | GO:0042476 | odontogenesis |
| 3 | GO:0022612 | gland morphogenesis |
| 4 | GO:0048705 | skeletal system morphogenesis |
| 5 | GO:0003007 | heart morphogenesis |
| 6 | GO:0090596 | sensory organ morphogenesis |
| 7 | GO:0048562 | embryonic organ morphogenesis |
| 8 | GO:1901135 | carbohydrate derivative metabolic process |
| 9 | GO:0006629 | lipid metabolic process |
| 10 | GO:0042592 | homeostatic process |
| 11 | GO:0008283 | cell proliferation |
| 12 | GO:0008219 | cell death |
| 13 | GO:0002376 | immune system process |
| 14 | GO:0030154 | cell differentiation |

| | | |
|---|---|---|
| 15 | GO:0051234 | establishment of localization |
| 16 | GO:0048731 | system development |
| 17 | GO:0019538 | protein metabolic process |
| 18 | GO:0016043 | cellular component organization |
| 19 | GO:0023052 | signaling |
| 20 | GO:0050896 | response to stimulus |
